# Electroporation-Based Gene Delivery and Whole-Organoid Imaging in Human Retinal Organoids

**DOI:** 10.1101/2025.06.07.658463

**Authors:** Keevon Flohr, Michael Janecek, Lingyun Wang, Vicente Valle, Shaohua Pi, Rui T. Peixoto, Susana da Silva

**Author notes:** co-correspondent authors.

## Abstract

Human retinal organoids (hRetOrg) derived from human induced pluripotent stem cells (hiPSCs) have emerged as powerful *in vitro* systems for studying retinal development, modeling retinal diseases, and evaluating therapeutic strategies. However, current genetic manipulation approaches, such as stable hiPSC line generation and viral transduction, are laborious, costly, and inefficient, with limited spatial specificity and high variability. Here, we report a rapid, scalable, and spatially precise electroporation-based platform for efficient plasmid-based gene delivery in early-stage hRetOrg. This method enables tunable and region-specific transfection of retinal progenitor cells without viral vectors or clonal selection. Coupled with resonant-scanning two-photon microscopy, this approach allows fast live cell imaging of whole organoids with subcellular resolution. This versatile system supports high-throughput genetic manipulation and imaging in intact hRetOrg, advancing studies of human retinal development, gene function, and disease.

**Motivation:** hRetOrgs offer an unprecedented platform for functional genetic studies of human retinal development and disease. However, existing methods for gene manipulation in hRetOrg are limited by low throughput, inefficiency, and lack of scalability, hindering systematic analysis of gene function and regulatory elements. To address these limitations, we developed a streamlined, high-efficiency pipeline that enables spatially targeted electroporation of hRetOrg during early retinogenesis, combined with fast, high-resolution imaging of whole organoids using two-photon microscopy, allowing studies at both tissue and subcellular scales.

## Introduction

Human retinal organoids (hRetOrgs), derived from embryonic stem cells (hESCs) or induced pluripotent stem cells (hiPSCs), are powerful *in vitro* model systems for studying retinal development, modeling inherited retinal diseases, and testing therapeutic strategies. They faithfully recapitulate key features of *in vivo* retinogenesis, including the temporal sequence of cell differentiation and the organized stratification of the neural retina [1-4]. Their ability to self-organize into laminated structures with a human-relevant cellular composition and organization makes them uniquely suited for investigating cell fate specification, tissue morphogenesis, and neural circuit formation. Despite these advantages, the limited availability of efficient and flexible tools for transgene expression remains a major barrier to fully harnessing the experimental potential of hRetOrgs systems.

Current approaches for genetic manipulations in hRetOrg, such as CRISPR/Cas9-based editing to generate isogenic stem cell lines, offer high precision and stable gene expression but require several steps of clonal selection, expansion, and validation that span several weeks, rendering them impractical for high-throughput applications. Additionally, each target gene or regulatory element requires the generation of a new edited line, limiting scalability and flexibility in experimental design. Viral transduction methods using adeno-associated virus (AAV) or lentivirus have seen only limited success in hRetOrg, with inconsistent efficiency and few reported implementations to date [5-8]. Viral approaches also present several drawbacks associated with limited packaging capacity, variability in transduction efficiency, and technical demands associated with producing quality high-titer viral stocks. A recent study raised additional concerns about viral-induced cell fate biases, reporting enhanced photoreceptor specification following early lentiviral transduction [7]. These findings underscore the need for caution when applying lentiviral methods in developmental studies. Moreover, most viral transduction protocols have been applied to late-stage organoids [9, 10], leaving early developmental stages largely unexplored.

To overcome these challenges, we optimized plasmid DNA electroporation as a non-viral gene delivery strategy for early stage hRetOrg. Electroporation transiently permeabilizes the cell membrane using brief electrical pulses, enabling rapid and cost-effective delivery of plasmid constructs without the need for clonal line generation or viral production. Importantly, electroporation can be timed to specific stages of organoid development, allowing targeted manipulation of distinct retinal progenitor populations. Spatial specificity can also be achieved by adjusting electrode placement, enabling localized gene delivery to defined organoid regions. These features establish electroporation as a powerful, high-throughput technique for spatially and temporally resolved gene manipulation during hRetOrg maturation.

Microscopic imaging is a critical tool in organoid research, enabling the visualization of structural organization, cellular dynamics, and functional processes. However, imaging intact organoids remains technically challenging due to their millimeter-scale size, dense cellular architecture, and optical heterogeneity. These features result in substantial light scattering and absorption, limiting imaging depth and resolution when using conventional one-photon microscopy. While tissue-clearing methods can improve light penetration and allow for whole-organoid imaging, they are incompatible with live imaging and thus preclude studies of dynamic processes. Two-photon scanning microscopy (TPSM) is well-suited for imaging whole organoids, particularly in live preparations[11]. By using near-infrared pulsed lasers and nonlinear excitation, TPSM confines fluorophore excitation to the focal plane, minimizing out-of-focus photobleaching and phototoxicity. This allows for deeper tissue penetration and improved signal-to-noise ratios compared to one-photon methods, while maintaining tissue viability during prolonged imaging. Recent integration of resonant galvanometer scanners has further enhanced imaging speed, enabling rapid full-frame acquisition. This is especially beneficial for capturing fast physiological events and for volumetric imaging of entire organoids, which would otherwise be limited by long acquisition times.

Here, we developed an integrated pipeline combining spatially targeted transfection of hRetOrg at early stages of retinogenesis via electroporation, followed by whole-organoid imaging of both tissue-cleared fixed and live imaging using mesoscopic two-photon microscopy. This approach supports rapid, high-throughput analysis of gene function and dynamic developmental processes across the whole organoid, opening new avenues for functional genomics, disease modeling, and circuit-level investigations in human retinal organoids.

## Results

### Electroporation of D42 hRetOrg with spatial control

hRetOrg used in this study were differentiated from a control hiPSC line based on previously established methods [1, 2, 12] (Figure 1A). To ensure consistency, we selected organoids that exhibited a well-defined retinal neuroblastic layer (NBL) spanning the entire structure. These high-quality hRetOrg were readily identifiable by the presence of a translucent, pseudo-stratified neural layer surrounding a central hollow, typically observed after day 30 (D30) of differentiation (representative examples in Figure 1A) [4]. Morphological selection criteria were validated by histological and molecular characterization of D45 organoids using the generic early retinal markers RAX and PAX6. We detected RAX and PAX6-positive cells forming a retinal neuroblastic layer throughout the entire organoid (Supplementary Figure S1), confirming its retinal identity. To test electroporation-based gene delivery in hRetOrg, we designed custom acrylic chambers incorporating a laser-cut central well measuring (6 × 6 × 2.5 mm), flanked by fixed stainless-steel electrodes positioned in parallel to generate a uniform electric field. This chamber design accommodates up to five D42 (week 6, W6) hRetOrgs, each approximately 750µm in diameter (Figure 1A), within a 50 µL volume, while maintaining consistent orientation across samples.

**Figure 1.**
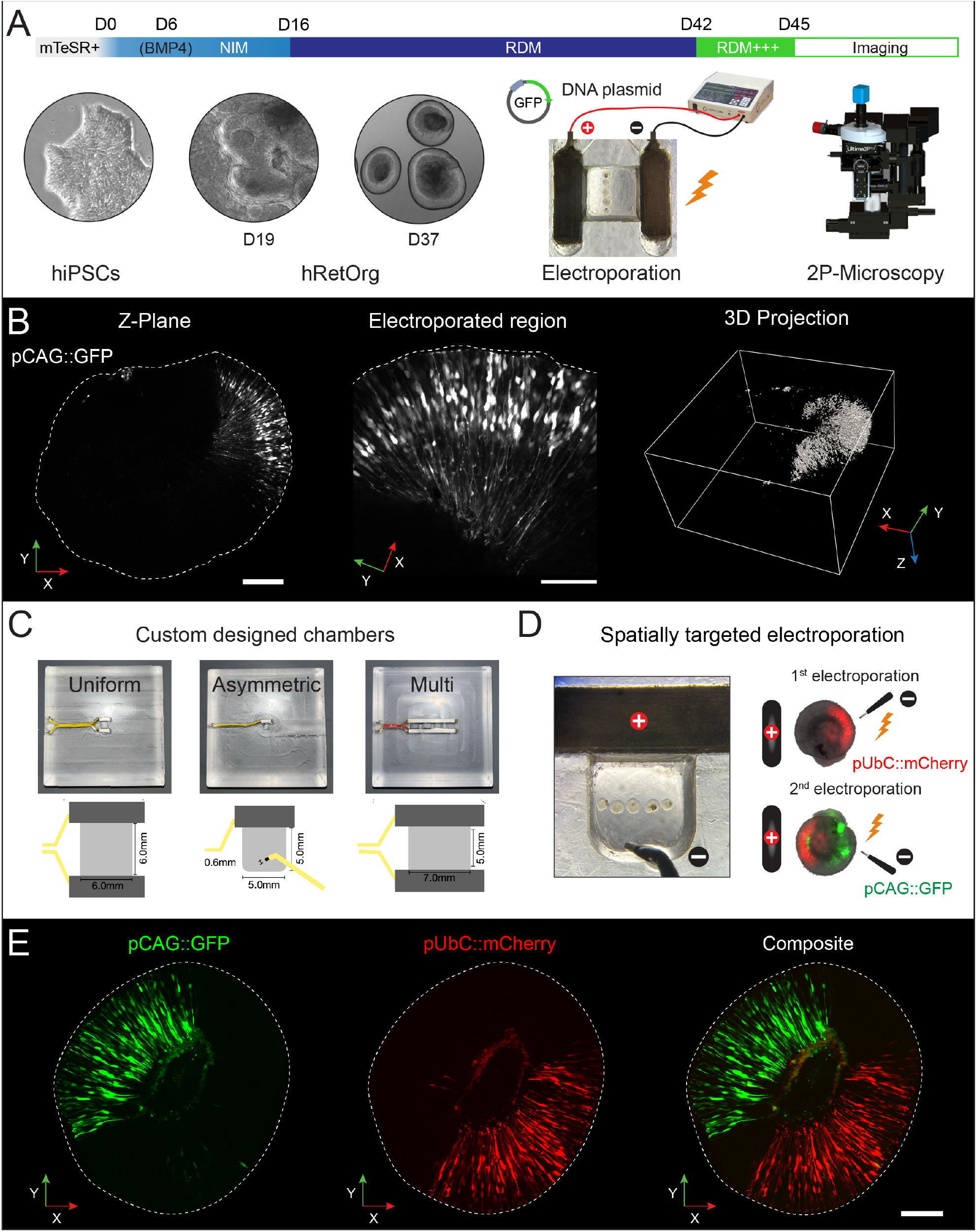
Spatially targeted electroporation in hRetOrg enables localized gene delivery. **(A)** Schematic of the experimental timeline and protocol. Human induced pluripotent stem cells (hiPSCs) were differentiated into retinal organoids (hRetOrgs) using a stepwise protocol involving neural induction medium (NIM) and retinal differentiation medium (RDM). Electroporation was performed at day 42 (D42), followed by imaging on day 45 using two-photon (2P) microscopy. Brightfield images show representative stages: undifferentiated hiPSCs, very early organoids at retinal cup stage (D19), and hRetOrgs at early retinogenesis period (D37). **(B)** Representative TPSM images of a D45 hRetOrg electroporated with pCAG::GFP at D42. Left: single Z-plane demonstrates spatial confinement of transgene expression, scale bar = 200 μm; Middle: higher magnification view of the electroporated region revealing radially aligned GFP-positive cells, scale bar = 100 μm; Right: 3D reconstruction of the entire organoid confirms focal transfection of a specific domain of the organoid. **(C)** Custom-designed electroporation chambers. Three chamber types were developed to enable different targeting strategies: uniform electric field (broad expression), asymmetric electrodes (focal targeting), and multi-slots (simultaneous transfections for higher throughput). Insets show the dimensions and electrode arrangements. **(D)** Image of the asymmetric chamber configured for spatially targeted electroporation. Diagram at right illustrates dual electroporation strategy: first with pUbC::mCherry, followed by a second electroporation with pCAG::GFP to target distinct organoid regions. **(E)** Representative whole-organoid 2P images of a Z plane showing spatially restricted expression of pUbC::mCherry (red) and pCAG::GFP (green), delivered via sequential electroporation. Composite image demonstrates non-overlapping, opposite region-specific expression patterns within the same organoid. Outlines mark the organoid boundary.

To assess transfection efficiency, we electroporated a DNA plasmid encoding enhanced green fluorescent protein (GFP) under the control of the ubiquitous CAG promoter at a final concentration of 150 ng/µL. Electroporation parameters were adapted from established ex vivo chick and mouse retina protocols [13-16] of five square-wave pulses at 40 V, each 50 ms in duration with 950 ms inter-pulse intervals. GFP expression was detectable approximately 8 hours post-electroporation using a fluorescence-equipped inverted digital microscope (EVOS 5000, Thermofisher) enabling rapid evaluation of transfection success.

To analyze the spatial pattern of GFP expression at high resolution, we imaged organoids three days post-electroporation (D45) using two-photon scanning microscopy (Figure 1A). Prior to imaging, organoids were fixed and tissue-cleared using a modified CUBIC protocol [17] to enhance optical transparency. Cleared samples were mounted between coverslip spacers to preserve their 3D morphology. Full-volume imaging of transfected RetOrg was performed using a two-photon scanning microscope (TPSM, Ultima 2P Plus, Bruker, WI) equipped with a 16X/0.8 NA long-working-distance objective (Nikon N16XLWD-PF) and 2-inch collection optics. This setup provides a large field of view (1.4 × 1.4 mm), well suited to the typical size of human retinal and brain organoids, enabling rapid scanning and seamless whole-organoid imaging without the need for tiling. GFP fluorescence was excited at 920 nm using a tunable laser (Insight X3, Spectra-Physics, CA) and imaged throughout the entire organoid volume.

Organoids were imaged in galvo-galvo mode with 2.67 sec/frame period and 2 μm z-spacing. Under these settings, an organoid with an average depth of 900 µm took approximately 20 min to image. (Figure 1B). GFP-positive cells were confined to a single domain of the organoid, oriented toward the anode, consistent with the directional nature of electroporation. Because GFP is a cytosolic reporter, it filled the entire cell, revealing the full morphology of transfected retinal cells. Most labeled cells extended from the apical to the basal surfaces of the retinal neuroblastic layer (NBL), a hallmark of retinal progenitor cells (RPCs) (Fig. 1B, left and middle panels). RPCs extend processes from the apical to the basal surfaces of the NBL and undergo interkinetic nuclear migration (INM) along this axis, depending on their cell cycle phase. This dynamic nuclear behavior has been shown to be recapitulated in hRetOrg [1]. Interestingly, the somas of most GFP-positive cells were clustered in a semi-discreet band of the NBL, suggesting that electroporated RPCs may be synchronized in both INM (Figure 1B, middle panel) and cell-cycle progression. Volumetric reconstruction revealed robust GFP expression localized to one region, confirming successful and spatially patterned gene delivery (Figure 1B, right panel).

To expand the versatility of our platform, we developed several custom electroporation chamber designs tailored to different experimental paradigms (Figure 1C). In addition to the standard dual fixed-electrode configuration that generates a uniform electric field (Figure 1C, left), we created a chamber with one fixed and one movable electrode. This asymmetric setup allows dynamic electrode positioning for more precise spatial targeting of electroporation (Figure 1C, middle). We also engineered a multi-slot chamber that enables high-throughput electroporation of up to four experimental conditions in parallel, facilitating the simultaneous transfection of multiple constructs under consistent field parameters (Figure 1C, right). To directly test whether our approach enables spatially defined gene expression, we performed sequential electroporations using the asymmetric chamber (Figure 1D). Each D42 hRetOrg underwent two rounds of electroporation with plasmids encoding complementary fluorescent reporter proteins expressed under ubiquitous promoters: pUbC::mCherry in the first round and pCAG::GFP in the second. To ensure robust expression, plasmid concentrations were increased to 600 ng/µL. Between electroporations, each organoid was rotated to target opposing domains, with the anode repositioned adjacent to the intended transfection region. Three days later, TPSM imaging revealed spatially distinct expression patterns of mCherry and GFP in opposite regions of the same hRetOrg (Figure 1E; Supplementary Movie S1), demonstrating effective spatial segregation of gene expression. Notably, organoids tolerated multiple rounds of electroporation without signs of structural damage, underscoring their robustness at this developmental stage. Together, these findings highlight the utility of our system for applications requiring fast and precise spatiotemporal control of gene expression in developing hRetOrg.

### Tunable electroporation settings for graded transfection efficiency

After establishing a baseline protocol for efficient gene delivery in hRetOrg, we assessed whether varying electroporation voltage could modulate transfection efficiency to accommodate different experimental needs. These include conditions requiring high transgene expression, as well as applications favoring sparse labeling, such as studies of cell-autonomous processes or long-term single-cell tracking via time-lapse imaging. To this end, we tested three voltages— 15 V, 40 V, and 65 V—while keeping the DNA plasmid concentration constant at 150 ng/µL. To account for variability in organoid morphology, size, and differentiation state, we selected high-quality D42 hRetOrgs ranging from 400–900 µm in diameter. For each voltage condition, at least eight organoids derived from independent differentiations were analyzed to ensure statistical robustness and reproducibility. Organoids were processed using the same tissue-clearing protocol as described previously (Figure 1). To assess the scalability of our imaging pipeline, we acquired full volumetric images of individual organoids using resonant-scanning mode of the TPSM. Imaging was performed at a ∼15 Hz frame rate with 16× frame averaging, resulting in an acquisition time of approximately 6–8 minutes per organoid, depending on imaging depth (Figure 2). This acquisition rate is compatible with large-scale studies, enabling a single system to image over 100 organoids per day.

**Figure 2.**
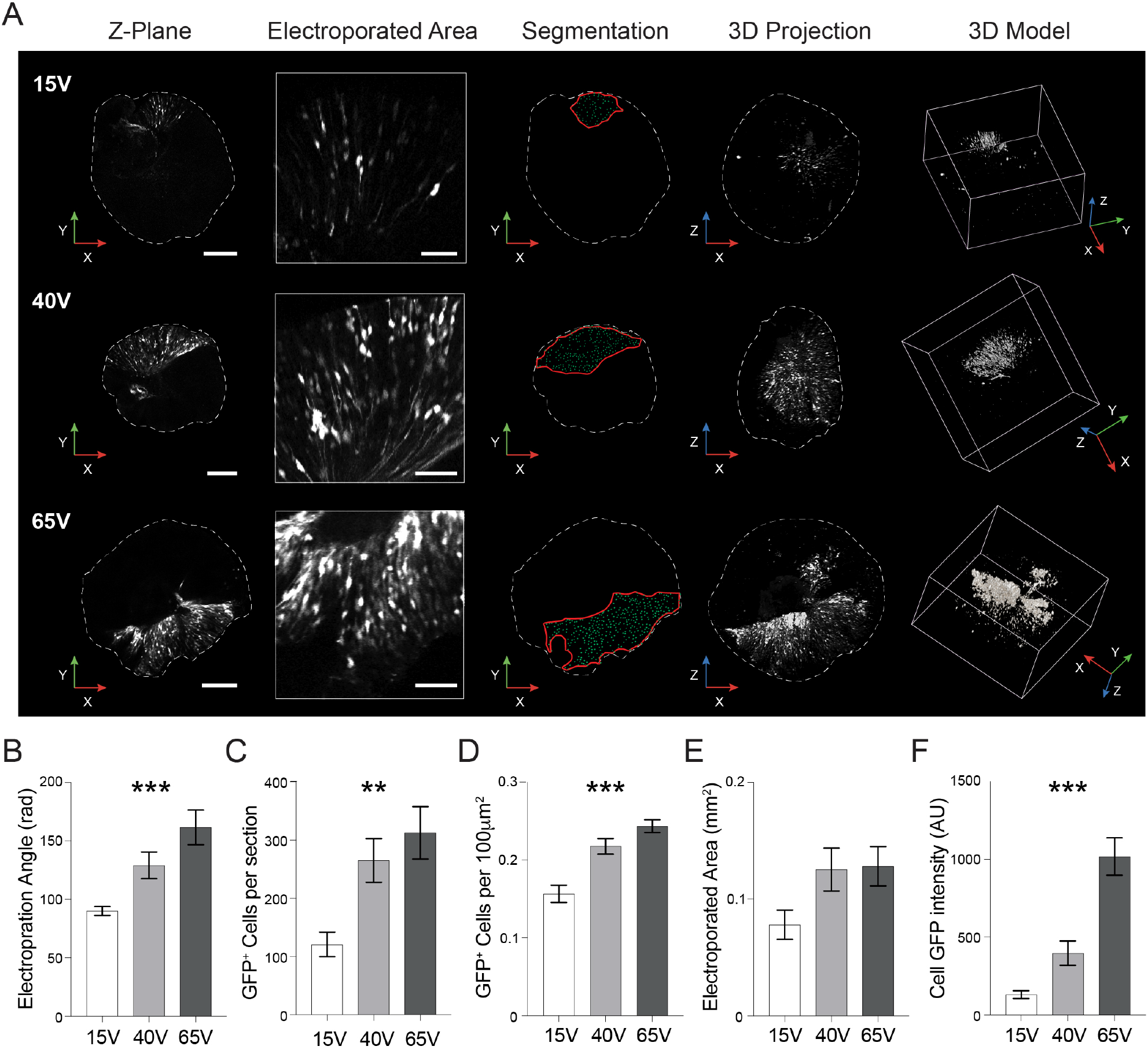
Electroporation voltage modulates the level and spatial extent of transgene expression in hRetOrg. **(A)** Representative 2P images showing the effect of increasing electroporation voltage (15V, 40V, 65V) on spatial targeting and transgene expression in D45 hRetOrgs electroporated with pCAG::GFP at D42. Columns (left to right): Z-plane projection of the whole organoid, scale bar = 200 μm; higher magnification of the electroporated region, scale bar = 50 μm; segmentation of GFP-positive cells within the organoid boundary (dashed line), with transfected regions highlighted (red outline); 3D projection of transfected cells in XZ view; and full 3D model showing spatial distribution and depth of gene delivery. **(B–E)** Quantification of electroporation outcomes across conditions, plots depict Mean ±SEM. Note an increase of all outcomes with voltage. **(B)** Angle of electroporated area (in radians). **(C)** Average of GFP-positive cells per section. **(D)** Density of GFP-positive cells within the targeted region. **(E)** Area of the electroporated region. **(F)** Mean GFP fluorescence intensity per cell. Mean Error bars represent standard error of the mean (SEM); n, number of organoids per condition = 8,9,9 for 15V, 40V and 65V, respectively. *, p < 0.01; ** p < 0.001; *** p < 0.001, one-way ANOVA.

To quantitatively assess electroporation efficiency, we performed automated cell segmentation on a stack of 11 consecutive optical sections centered around the plane with the highest density of electroporated cells (Figure 2A). From this region, we quantified several key metrics: 1) the average angle of electroporated region relative to the geometric center of the hRetOrg, 2) the total number of GFP-positive cells, 3) the average density of GFP-positive cells, 4) the area of electroporated region and 5) the average GFP signal intensity per cell. This targeted analysis approach was chosen to optimize both spatial resolution and segmentation performance of GFP-positive cells. Focusing on a central, high-signal z-stack allowed us to capture representative electroporation patterns while ensuring accurate and consistent cell detection across samples. This method provided a reliable and scalable framework for quantifying transfection outcomes across different experimental conditions.

We observed a clear voltage-dependent increase in both electroporation efficiency and the spatial extent of transgene expression. The average angle of the electroporated region expanded with increasing voltage (Figure 2B, Mean ± SEM; 15V: 90.0°± 3.9°; 40V: 129.1°± 11.4°; 65V: 161.6°± 14.9°; one-way ANOVA, *p* < 0.001; *n* = 8, 9, and 9).

Correspondingly, the number of GFP-positive cells in the segmented 2D optical section increased with voltage (Figure 2C, Mean ± SEM; 15V: 120.8 ± 20.9; 40V: 268.5 ± 37.5; 65V: 312.4 ± 45; one-way ANOVA, *p* = 0.004) with respective increase in cell density when normalized to electroporated area (Figure 2D, Mean ± SEM; 15V: 0.16 ± 0.01; 40V: 0.22 ± 0.01, and 0.24 ± 0.01 cells/100μm^2^; one-way ANOVA, *p* < 0.001) that showed a modest but not statistically significant increase with voltage (Figure 2E, Mean ± SEM; 15V: 0.078 ± 0.013 mm^2^; 40V: 0.125 ± 0.018 mm^2^; 65V: 0.128 ± 0.016 mm^2^; one-way ANOVA, *p* = 0.07). The fluorescence signal intensity per cell also increased significantly, indicating that electroporation voltage can be leveraged to control overall construct expression level (Figure 2F, Mean ± SEM; 15V: 131.1 ± 25; 40V: 397.6 ± 78; 65V: 1017.6 ± 120.1 arbitrary units; one-way ANOVA, p < 0.001). These findings demonstrate a positive correlation between electroporation voltage and transfection efficiency, both in terms of expression level and spatial coverage. Notably, the 15 V condition produced a small circular, spatially confined population of labeled cells, which may be particularly useful for experiments requiring sparse, localized gene expression. These trends are visually represented in the 3D projections shown in Figure 2A and the accompanying supplementary videos (Supplementary Movies S2, S3 and S4).

### Characterization of electroporated cells in D45 hRetOrg

D42∼D45 in hRetOrg development corresponds to an early stage of retinogenesis, during which the organoids are predominantly composed of multipotent RPCs undergoing cell divisions to generate more RPCs, while some RPCs exit the cell cycle to give rise to early born retinal neurons. Retinogenesis follows a well-characterized temporal order of neurogenesis, with retinal ganglion cells (RGCs) being the first to become postmitotic, followed by cone photoreceptors, and subsequently horizontal and amacrine cells (H/ACs) [18]. Therefore, transfection of hRetOrgs with a plasmid encoding a transgene under the control of a ubiquitous promoter is expected to result in labeling of all these retinal cell types in proportions that reflect their developmental timing. To confirm this developmental cellular composition, we performed immunostaining on cryosections of D45 organoids that had been electroporated at D42 with pCAG::GFP. We used well-established molecular markers specific to the expected retinal cell types at this stage, specifically RAX (labeling most retinal cells excluding first-born RGCs and first-born H/ACs), CHX10 (also known as VSX2, labeling RPCs), ISLET1/2 (labeling RGCs (ISL1) and cones (ISL2)), OTX2 (labeling cones and a subset of progenitors transitioning to cones and horizontal cells), and TFAP2α (labeling H/ACs).

Analysis of organoid cross-sections revealed that electroporation was spatially restricted to a defined area, consistent with the localized GFP signal observed in whole-organoid imaging (Figures 1 and 2). Similarly, examination on hRetOrg cryosections showed that the majority of GFP-positive cells displayed characteristic morphological features of RPCs, with radial processes extending from the apical to basal surfaces of the NBL (Figure 3A, right panel).

**Figure 3.**
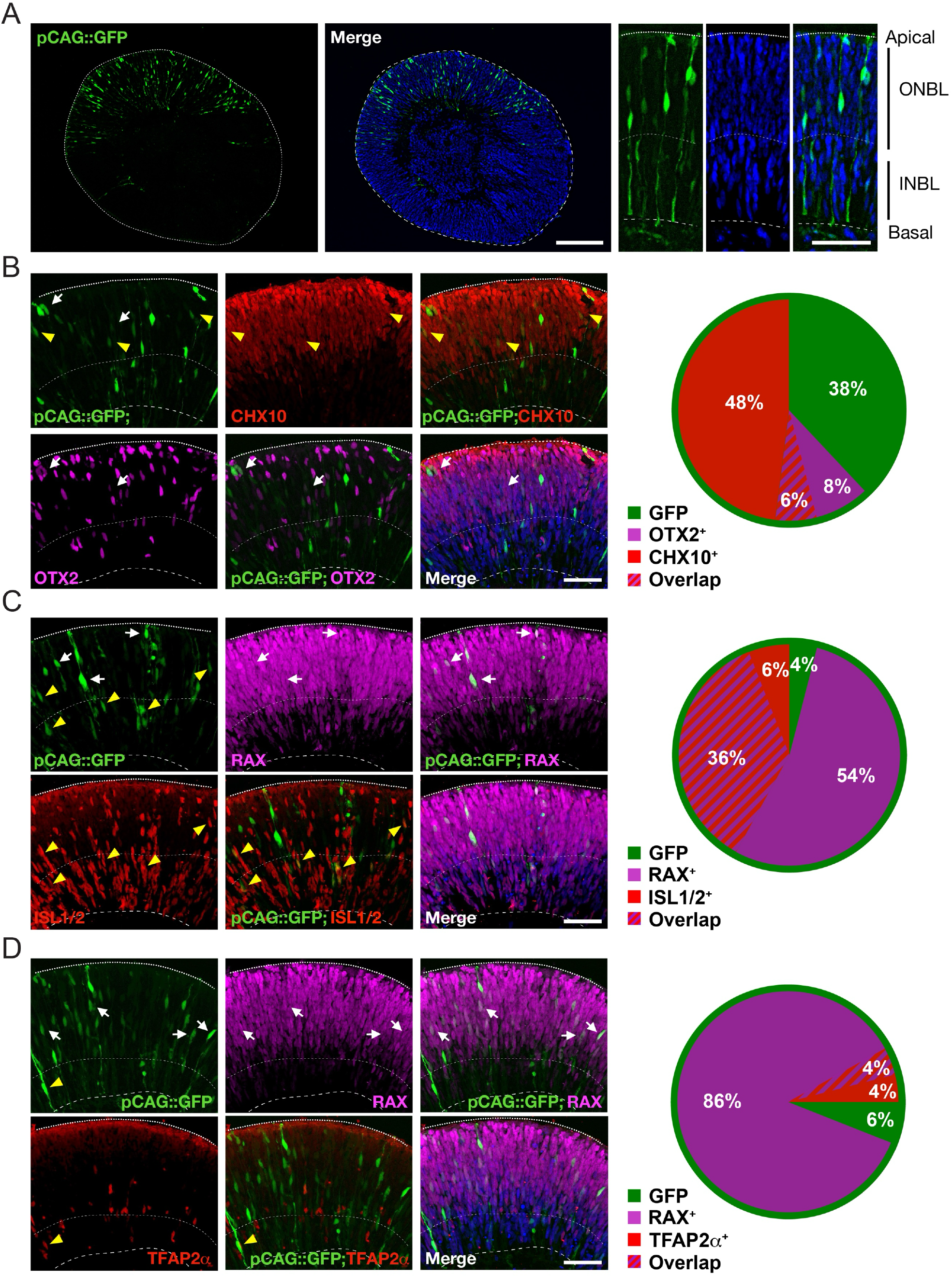
Electroporation at early stages of hRetOrg development predominantly targets retinal progenitor cells. **(A)** Immunohistochemical analysis of a cross-section of a D45 hRetOrg electroporated with pCAG::GFP at D42. Left and middle panels: whole-organoid views showing GFP-positive cells confined to one domain of the organoid, Hoeschst counterstained. Right panels: higher magnification images showing GFP-positive cells aligned radially from the apical to basal surface of the neuroblastic layer (NBL), consistent with RPC morphology. **(B)** Confocal images of organoid sections stained for the retinal cell markers CHX10 (also known as VSX2) and OTX2, counterstained with Hoechst. GFP-positive cells co-localize with CHX10 (red, yellow arrowheads) and OTX2 (magenta, white arrows). Pie chart quantifies marker expression among GFP-positive cells: 48% GFP^+^/CHX10^+^, 8% GFP^+^/OTX2^+^, 6% GFP^+^/CHX10^+^/OTX2^+^, and 38% GFP only. **(C)** Additional immunostaining for the early retinal markers RAX and ISLET1/2. Many GFP-positive cells co-express RAX (purple, white arrows), with a subset also expressing ISLET1/2 (red, yellow arrowheads). Pie chart quantifies marker expression among GFP-positive cells: 90% GFP^+^/RAX^+^, 42% GFP^+^/ISLET1/2^+^, 36% GFP^+^/ RAX^+^/ ISLET1/2^+^ and 4% GFP only. **(D)** Co-labeling with RAX and TFAP2α identifies few GFP-positive cells co-expressing TFAP2α (red, yellow arrowheads). The majority of GFP-labeled cells co-express RAX (purple, white arrows). Pie chart quantifies marker expression among GFP-positive cells: 86% GFP^+^/RAX^+^, 4% GFP^+^/TFAP2α^+^, 4% GFP^+^/RAX^+^/TFAP2α^+^, and 6% GFP^+^ only. Dotted lines in all panels indicate boundaries of the organoids: apical, basal and approximate boundary between the outer neuroblastic layer (ONBL) and inner neuroblastic layer (INBL). Scale bars: 200 μm, left and middle panels in A, and 50 μm in all other panels.

Quantification of marker co-localization with GFP-positive cells demonstrated that, on average, 54% of electroporated GFP-positive cells co-expressed CHX10 (also known as VSX2, RPC marker) (Mean ± SEM; 54% ± 3%, *n* = 5), while 14% of those co-localized with OTX2 (mostly cones) (14% ± 2%, *n*=5) (Figure 3B), with 6% of GFP-positive cells staining for both markers (RPCs about to become a cone or horizontal cell) (6% ± 3%, *n*=5). Co-localization analysis of GFP-positive cells with RAX and ISLET1/2, demonstrated that on average 90% of electroporated GFP-positive cells co-localized with RAX (all retinal cells except first-born RGCs and first-born H/ACs) (90% ± 1%, *n*=5) and 42% co-labeled with ISL1/2 (42% ± 2%, *n*=5) (Figure 3C). ISL1/2 antibody labels both ISL1 (RGCs) and ISL2 (cones) proteins. Cones and RGCs occupy a distinct lamina in the retina, with cone photoreceptors located to the apical surface and RGCs on the opposed basal surface of the NBL, forming what has been initially named inner neuroblastic layer (INBL). Based on the laminar location of ISL1/2-positive cells, we estimate that from the GFP-positive and ISL1/2-positive population, about 25% are likely RGCs (co-labeled cells located in the INBL, 25% ± 3%) while 17% are cones (co-labeled cells in the ONBL, 17% ± 4%). In this immunostaining combination, 36% of GFP-positive cells were labeled for both RAX and ISL1/2, likely corresponding to recently born RGCs and cones (36% ± 2%, *n*=5). In a separate round of immunostaining, we found that 90% of GFP-positive cells co-localize with RAX (90% ± 2%, *n*=5), while 8% were TFAP2α-positive (H/ACs) (8% ± 1%, *n*=5) with 4% being triple-labeled, likely corresponding to recently specified H/ACs (4% ± 2%, *n*=5) (Figure 3D). Altogether, these quantitative analyses confirm the expected type and ratio of retinal cells present in the organoid at this early stage of retinogenesis, suggesting that electroporation did not influence cell fate specification in hRetOrg.

### Live two-photon imaging of electroporated hRetOrgs enables time-lapse studies with sub-cellular resolution

To further demonstrate the potential of our electroporation and imaging pipelines, we applied resonant-scanning two-photon microscopy for live, time-lapse imaging of electroporated hRetOrgs. This approach enabled high-resolution visualization across spatial scales, from whole-organoid cell migration to subcellular organelle trafficking, without requiring tissue clearing. Time-lapse imaging of cell migration was performed on D45 hRetOrgs electroporated at D42 with pCAG::GFP. High-speed resonant scanning allowed tracking of individual GFP-labeled retinal cells at 5-minute intervals over a 45-minute period (Figure 4A). The resulting time series captured the progressive nuclear translocation of an RPC undergoing typical INM behavior along the apical-basal axis of the NBL. Yellow arrowheads indicate the nucleus position over time, demonstrating the resolution of our system for visualizing dynamic cell behavior in live whole organoids. We also performed real time subcellular imaging by expressing a mitochondrially localized GFP expressed under a ubiquitous promoter using pEF1α::mitoGFP, at a final concentration of 600 ng/µl [19]. Electroporation of this construct into D42 hRetOrgs enabled visualization of mitochondrial trafficking along the apical-basal axis of retinal cells (Figure 4B). By restricting imaging to 50 μm depth we could acquire z-stacks at 30-second intervals, allowing continuous tracking of individual mitochondria. Together, these findings establish the feasibility of using resonant-scanning two-photon microscopy for high-resolution, multiscale imaging in electroporated hRetOrgs. Our approach enables longitudinal tracking of cellular and organelle dynamics in live, non-cleared organoids, offering a powerful platform for investigating cellular processes in real time.

**Figure 4.**
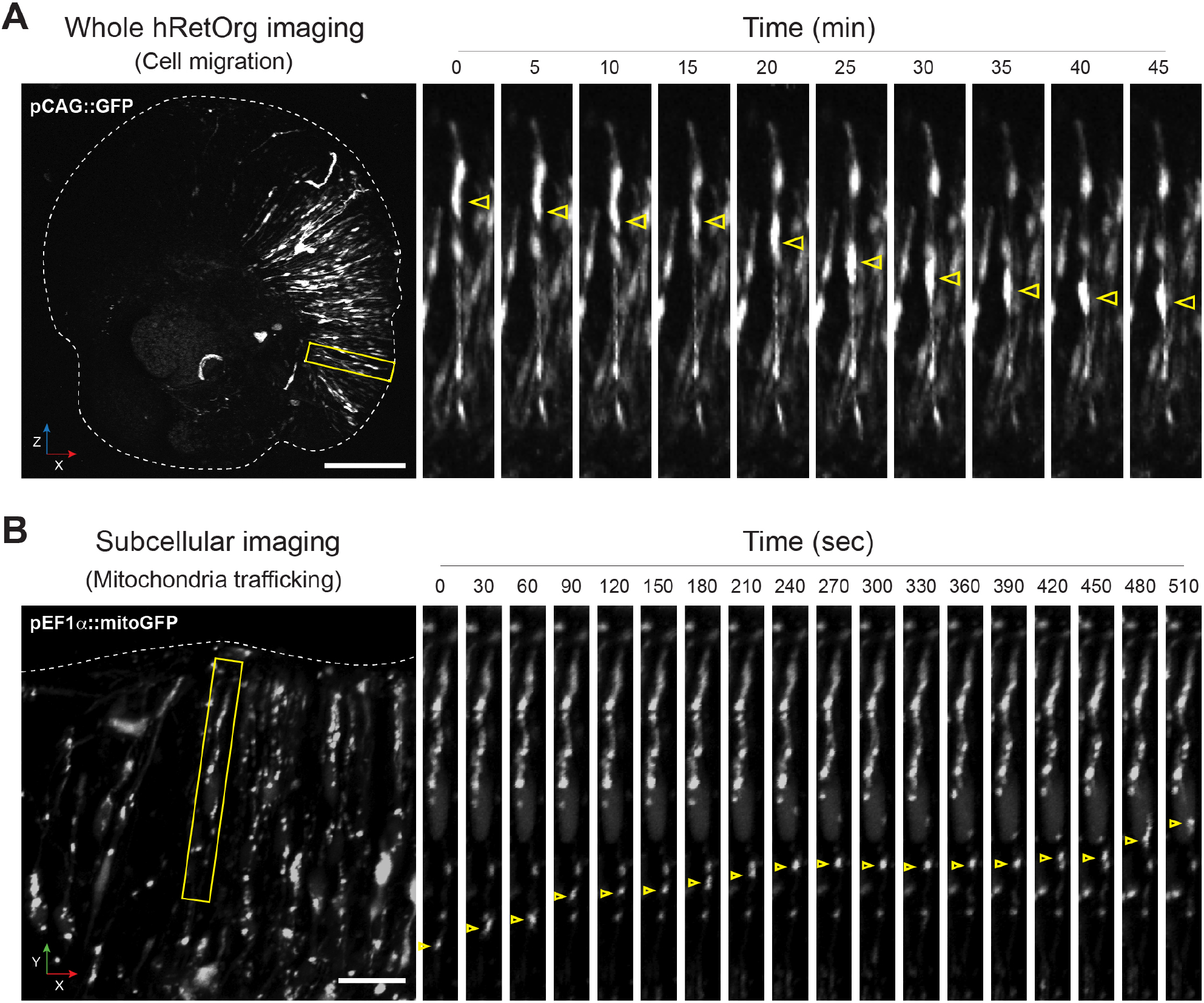
Two-photon live imaging reveals dynamic cell and organelle behaviors in whole hRetOrg. **(A)** Time-lapse two-photon imaging of a whole D45 hRetOrg electroporated with pCAG::GFP at D42 highlighting interkinetic nuclear migration. Left: Z-projection of the entire organoid showing region of interest (yellow box) selected for time-lapse imaging, scale bar = 200 μm. Right: sequential frames from the boxed region acquired at 5-minute intervals over 45 minutes. Yellow arrowheads track the position of a single migrating GFP-positive cell soma, demonstrating movement along the apicobasal axis of the neuroblastic layer. **(B)** High-resolution subcellular imaging of mitochondrial transport in hRetOrgs electroporated with pEF1α::mitoGFP. Left: representative image showing the region selected for time-lapse imaging (yellow box). Right: sequential frames captured at 30-second intervals over 510 seconds. Yellow arrowheads follow a single GFP-labeled mitochondrion, illustrating directed intracellular trafficking along the apicobasal axis, scale bar = 50 μm.

### Limitations of the Study

Electroporation will only target cells undergoing cell division and therefore this method is most effective in early-stage organoids which are enriched with RPCs; older organoids may require optimized parameters to balance transfection efficiency and cell viability. Moreover, this study is based on episomal plasmid DNA, which enables efficient gene delivery but may result in transient transgene expression due to plasmid dilution in highly proliferative RPCs. To overcome this limitation, genomically integrative strategies can also be delivered via plasmid, such as transposon-based systems. Due to challenges in segmenting highly elongated RPCs labeled with cytoplasmic markers in 3D datasets, this study relied on 2D optical sections for quantitative analysis. Incorporating reporter genes with nuclear localization signals in future experiments could significantly improve 3D volumetric segmentation and analysis accuracy.

Electroporation inherently produces mosaic expression, which may limit certain applications but can be advantageous for comparing neighboring transfected and untransfected cells within the same tissue. For uniform expression across all cells, generating stable hiPSC lines may be more suitable.

## Discussion

In this study, we present a robust, scalable, and versatile platform for gene delivery in human retinal organoids (hRetOrg) using electroporation, integrated with high-resolution whole-organoid imaging. Our results establish electroporation as a rapid and efficient method for transfecting early-stage hRetOrg, overcoming key limitations of traditional genetic manipulation approaches, such as the extended timelines, cost, and labor associated with generating isogenic hiPSC lines, and the expression variability and cargo constraints of viral vectors.

Electroporation is highly efficient and reproducible in hRetOrg because the apical side of the retinoblast, where RPCs undergo mitosis and are thus more susceptible to plasmid entry into the nucleus, faces outward and directly contacts the DNA plasmid mixture. Applying electroporation to other types of organoid tissues will likely require optimization of both the electroporation parameters and the site of plasmid delivery. In hRetOrg, we show that electroporation can be precisely timed to target RPCs within a defined developmental window. When applied at early stages of organoid differentiation, electroporation labels RPCs and their early-born progeny, including RGCs, cone photoreceptors, and H/ACs. This developmental specificity, combined with localized spatial control, enables targeted perturbation of discrete cell populations, supporting mechanistic studies of lineage specification, neurogenesis, and retinal patterning in a human-relevant model.

We further show that transfection efficiency and distribution can be finely tuned by modulating electroporation voltage, accommodating experimental designs ranging from sparse single-cell labeling to broad regional transfection. To enhance precision and throughput, we engineered custom electroporation chambers, including movable-electrode and multi-slot configurations, enabling spatial targeting and parallel processing of multiple conditions. Sequential electroporation with distinct plasmids enabled combinatorial labeling of organoid regions, facilitating dissection of spatially restricted gene functions and interregional interactions. To visualize developmental processes across scales, we integrated this gene delivery method with mesoscopic two-photon microscopy for both fixed and live organoids. In cleared tissues, full-volume imaging captured patterned gene expression with high resolution. In live organoids, we recorded dynamic events such as interkinetic nuclear migration and mitochondrial trafficking, enabling longitudinal tracking of cellular and subcellular processes. Both galvo-galvo and resonant scanning modes proved compatible with our system, maximizing high-resolution or high-throughput as needed. A galvo-resonant scanner provides frame rates up to 30 Hz, allowing for rapid volumetric imaging with a temporal resolution sufficient to capture dynamic cellular activity. With high fluorescence signals, this configuration enables full organoid volumes to be imaged in as little as 2–3 minutes, making the platform highly compatible with high-throughput applications such as drug screening or phenotypic assays in disease models.

Importantly, the ability to image live, uncleared organoids opens new opportunities for continuous, longitudinal studies of developmental trajectories, cellular behavior, and responses to environmental or genetic perturbations. Additionally, we developed a quantitative segmentation pipeline for systematic assessment of electroporation angle, transfected cell counts, and fluorescence intensity, enabling standardized, reproducible analysis across experimental conditions. In sum, our platform provides a fast, cost-effective, and spatially precise alternative to viral transduction and stable hiPSC editing for genetic manipulation in hRetOrg. By enabling efficient gene delivery, multiscale imaging, and quantitative analysis, this approach significantly expands the experimental toolkit for retinal organoid research and opens new avenues for functional screening, spatial gene perturbation, and developmental modeling. Moreover, this approach provides a broadly adaptable framework for genetic manipulation and live imaging in organoid models of other human tissues, with relevance to both basic developmental biology and translational research.

## Methods

### Human induced pluripotent stem cells (hiPSC) maintenance

Cells were cultured in mTeSR™ Plus medium (STEMCELL Technologies, #100-0276) at 37 °C with 5% CO_2_ in a humidified incubator. Cultures were maintained in 6-well plates (Corning, #3516) pre-coated with hESC-qualified Matrigel (Corning, #354277), following WiCell-recommended protocols. Cells were passed at approximately 80% confluency every 4–5 days.

### Differentiation of human Retinal Organoids (hRetOrg)

The differentiation of hRetOrg was performed based on previously described protocols [Zhong et al., 2014 Cowan et al. Harkin]. Briefly, hiPSC were dissociated into single cells using Accutase (Thermofisher Scientific, # 00-4555-56) and mechanical trituration with a P1000 pipette in 1ml of mTeSR™ plus medium containing 10µM Rock inhibitor Y-27632 (STEMCELL Technologies, #72304). Cells were then plated onto 100mm petri dishes (VWR # 25384-088) in10ml of mTeSR™ plus to promote embryoid bodies (EBs) formation. On Day (D) 1, approximately 1/3 of the medium was exchanged for neural induction medium (NIM) containing DMEM/F12 (Gibco, #11330057), 1% N2 supplement (Gibco, #17502048), 1x NEAAs (Sigma, #M7145), and 2mg/ml heparin (Sigma, # H3149). On D2, approximately 1/2 of the medium was exchanged for NIM. On D3 EBs were plated in new 60mm petri dishes (Corning #430166) in NIM. Half of the medium was changed every day until D6. At D6, BMP4 (R&D, #314-BP) was added to the culture at a final concentration of 1.5nM. At D7, EBs were transferred to 60mm dishes previously coated with Growth Factor-Reduced (GFR) Matrigel (Corning, #356230) and maintained with daily NIM changes until D15. On D16, NIM was exchanged for retinal differentiation medium (RDM) containing DMEM (Gibco, #15140122) and DMEM/F12 (1:1), 2% B27 supplement (without vitamin A, Gibco#12587-010), NEAAs, and 1% penicillin/streptomycin (Gibco, #15140122). Feeding was done daily until D27. On D28, EBs were dislodged from the plate by checkerboard scraping, using a 10µl or 200µl pipette tip. Aggregates were washed three times and then maintained in 6 well culture plate (Greiner bio-one, #657185) in RDM medium. The medium was changed every 2-3 days until D41. During this time, hRetOrg were sorted based on their morphology under a stereoscope with transmitted light. hRetOrg were clearly identified based on the presence of a typical phase-bright, pseudo-stratified neuroblastic epithelium, and an inner hollow. On D42, the medium was changed to RDM plus, which consisted of RDM supplemented with 10% FBS (Gibco, #16140071), 100µM Taurine (Sigma, #T0625) and 2mM GlutaMax (Gibco, #35050061). Feeding was done every 2-3 days, depending on the color of the medium.

### Electroporation of hRetOrg

D42 hRetOrg presenting typical neuroblastic-like retinal morphology in at least 80% of the entire organoids were electroporated with DNA plasmids prepared using Qiagen endo-free Maxi kits The plasmids utilized were pCAG::GFP (a gift from Connie Cepko, addgene #11150), pUbC::mCh (a gift from Connie Cepko) or pEF1α::Mito-cGFP (addgene #188904). Final concentration of each plasmid varied between experiments, ranging from 150 ng/µl to 600 ng/µl, as detailed in the text. pEF1α::mito-cGFP (600ng/µl) was mixed with pUbC::mCh (150ng/µl), which was utilized as a co-electroporation marker. Purified DNA plasmids were diluted in 1X PBS to a total volume of 50µl, when using the uniform or multi-slot electroporation chambers, or 40µl, for the non-uniform electric field chamber (specific dimensions of each chamber in Fig. 1C), and loaded in the chamber. Organoids were then transferred one by one using wide-bore tips (Thermo Scientific™ 9405123) to the electroporation chamber under a dissection scope inside a biosafety cabinet. Each chamber could fit on average five D42 organoids, of ∼ 750µm in diameter). Electroporations were conducted with a NEPA21 type II Nepagene electroporator and 5 pulses of variable voltage (either 15V, 40C or 65V) of 50msec pulse duration with 950msec interpulse interval were applied. After electroporation organoids were transferred using wide bore tips to a tissue culture plate with RDM plus medium and finally transferred to a new plate and returned to the incubator for additional 3 days. While in culture, organoids were imaged under EVOS M5000 (Thermofisher) equipped with fluorescence filters for GFP and TX Red to check for electroporation efficiency.

### Two photon imaging

For imaging of fixed organoids, D45 hRetOrg electroporated at D42 were fixed with 4% PFA for 30min at RT, washed with 1X PBS and subjected to a CUBIC-based tissue clearing protocol as described in Muntifering et al., 2018, for approximately 1 week until fully transparent. Organoids were then mounted in R2 solution using coverslip spacers (SS1×9-Secure Seal, #654002, Grace Bio-Labs). Imaging was performed using a scanning two-photon excitation fluorescence microscope (Ultima 2P Plus, Bruker, WI). The microscope was controlled using PrairieView software control (vX5.5, Bruker, WI), a two-photon excitation was provided by a tunable femtosecond pulsed infrared laser (Insight X3, Spectra-Physics, CA), set to 920 nm for optimal excitation of GFP and 1100 nm for mCherry. The laser beam was directed into the microscope via a dichroic mirror (ZT1040dcrb-UF3, Chroma, VT). High-resolution images (Figure 1) were acquired in galvo-galvo mode at a resolution of 1024 × 1024 pixels through a 16× water-immersion objective (Nikon 16×/0.8 NA) in overscan mode (zoom at 0.8x) resulting in a 1.42×1.42mm field of view (1.34 µm/pixel). Volumetric datasets of each organoid were acquired by collecting sequential optical sections over a total imaging depth of 750-900 µm with 1 µm axial (z) steps and scanned with a dwell time of 6.2 µs/pixel and photomultiplier tube (PMT) gain settings of 600 HV. For comparative experiments analyzing electroporation voltage settings (Figure 2), imaging was performed using resonant-galvo scanning to increase throughput.

Images were acquired at ∼15.2 Hz with 16 frame averages and 2 µm z-step increments. Live-cell imaging of hRetOrg was performed in a 3.5 mm Petri dish modified with a custom polypropylene cup (2 mm in diameter) affixed to the bottom. Imaging medium consisted of AMES solution supplemented with 50 mM HEPES. The culture medium was maintained at 36.5 °C using a culture dish microincubator (Warner Instruments DH-40iL) regulated by a temperature controller (Warner Instruments TC-324C). Whole-organoid live-cell imaging was performed in resonant-galvo mode, with images acquired at 3 µm z-steps and a frame interval of 5 minutes. For mitochondrial imaging, a 50 µm section containing the neuroblastic region was imaged in resonant scanning mode at a frame rate of 30 seconds per frame. All z-stacks were acquired sequentially and subsequently processed for three-dimensional visualization and quantitative analysis in ImageJ.

### Two-photon microscopy Imaging Analysis

Cell segmentation was performed using a customized pipeline developed in MATLAB (MathWorks, USA). Specifically, the contrast of the acquired two-photon microscopy images was first enhanced by gamma correction and saturation adjustment to highlight the organoid tissue and electroporated regions. Images were then binarized by global threshold, and morphological closing and hole-filling were applied to segment the organoid tissue. The area was further smoothed using morphological opening operations, Gaussian filtering, and subsequent re-binarization to refine the organoid segmentation. Subsequently, to identify the electroporated regions, local intensity maxima were conducted in the segmented organoid regions. Pairwise distance between the detected maxima was computed to connect adjacent points by a pre-defined threshold, generating an initial mask for the electroporated regions. Finally, another set of morphological closing and opening operations were applied to refine the segmentation of electroporated regions. The process was iterated for all depth slices. For each depth slice, retinal cells were detected as local intensity maxima within the electroporated regions. To calculate the signal intensity of each retinal cell, a circular region with radius (3 pixels) centered at each detected cell location was determined and the pixel intensities were averaged. Additionally, the angle of the electroporated region was measured manually in ImageJ (NIH, USA). For each organoid, we quantified their tissue and electroporated regions in 3-D using volume size as readouts. Then, we identified the depth slice with the highest detected electroporated cell count and selected a stack of 11 slices—including the reference slice plus five equally spaced slices above and five equally spaced slices below at intervals of one slice— to calculate retinal cell counts, signal intensity, and electroporated region angle. The values from all 11 slices were averaged to derive the representative metrics for each organoid. The reason we performed the analysis of electroporated cells counts in 2-D only was that the cells, mostly retinal progenitors, are tightly arranged, making it extremely hard to isolate them in the 3-D space in the high-speed resonant imaging mode. We selected 11 slices rather than averaging across the entire volume depth to avoid underrepresenting cell counts and electroporated regions, as edge slices of the volume often contain significantly fewer electroporated regions and cells. By focusing on the central region around the slice with the highest cell count, we ensure that our quantification is both representative and minimizes potential biases introduced by sparsely populated peripheral regions. For volumetric rendering of organoids, images were downsampled to 8-bit in ImageJ and 3D-projected using Y-axis rotation with brightest point projection. The lower transparency threshold was set to 5, and interior depth cueing was adjusted to 40%. Final 3D models were generated in 3D Slicer using the volume rendering plug-in.

### Immunostaining of hRetOrg cryo-sections

Electroporated hRetOrg were fixed with 4% PFA at RT for 30 min, washed 3 times with PBS, and cryoprotected in 1x PBS-30% sucrose at 4°C O/N. Next day, organoids were embedded in a mixture of OCT and PBS-30% sucrose (1:1) in individual blocks in the same mixture in dry-iced 90% ethanol. Cryosections of 16µm thickness were alternated across Superfrost Plus slides (Fisher, #12-550-15). Once fully dried, slides were stored at -80°C until further processing. On the day of staining, slides were removed from -80°C and allowed to warm up to RT for approximately 20 min. Slides were washed 3 times with PBS, and blocked 1h at RT with 5% Donkey Serum and 0.1% Triton X-100 in 1x PBS. Primary antibodies were diluted in 2% Donkey Serum in 1X PBS and incubated at 4°C O/N. Primary antibodies used included anti-Rx (Takara, #M229, 1:1000 dilution); anti-VSX2/CHX10 (Santa Cruz, #sc-365519, 1:100 dilution); anti-OTX2 (R&D, #AF1979SP, 1:50 dilution); anti-TFAP2α, DSHB, clone 3B5, 1:35 dilution); and anti-Islet1/2 (DSHB, clone 39.4D5, 1:200 dilution). Slides were rinsed 3 times, 5 min each, with 1x PBS and incubated with appropriate donkey raised Alexa-Fluo secondary antibodies (Thermofisher) for 1 hour at RT and counterstained with Hoechst (Thermofisher, catalog #62249). Slides were then mounted with Fluoromount-G solution (Southern Biotech, #0100-01) and imaged using an Olympus fv 1200 Confocal with 40x, NA1.3 objective.

### Co-localization analysis

For the quantification of co-localizations of electroporated GFP-positive cells with specific retinal cell type markers, namely ISL1/2, OTX2, RAX, CHX10 and TFAP2α, z-stacks images of immunostained organoid cryosections were obtained using the Leica Thunder System, followed by maximum projections. Cell counting was then performed using Fiji (version 2.14.0), first by drawing a 200 per 200 μm region that would encompass the electroporated patch, followed by manual counts by using the Cell Counter plugin. GFP-positive cells were determined to be positive when co-labeling with Hoechst (nuclear marker). Percentages of the different cell types being double labeled with GFP positive cells were then averaged from all of the organoids analyzed (n=5).

## Supplementary Material

**Supplementary Figure S1.**
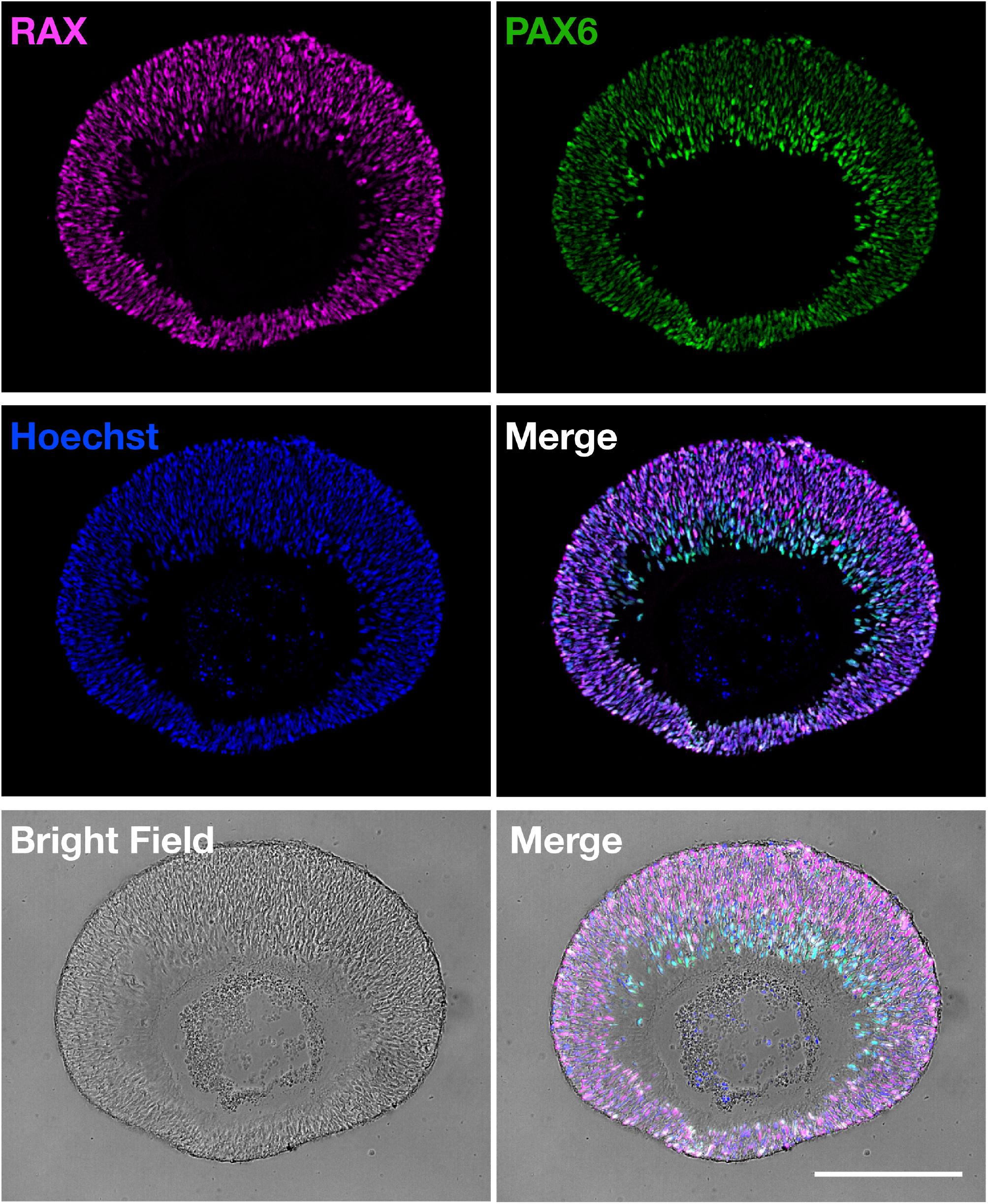
Characterization of retinal identity on a D45 hRetOrg. Cryosection of a representative 45 hRetOrg immunostained for the early retinal progenitor markers RAX (magenta) and PAX6 (green), counterstained with Hoechst (blue) to label nuclei. Both RAX and PAX6 are broadly expressed throughout the neuroepithelial layer, confirming the retinal identity. The merged fluorescence image shows co-expression and spatial overlap of these markers. Brightfield and corresponding overlay images demonstrate the pseudostratified organization of the retinal neuroblastic layer. Scale bar: 200 μm.

**Supplementary Movie S1. Opposed Spatially targeted domains of electroporation in early hRetOrg. Sequential electroporation of pCAG**

**Supplementary Movie S2. D45 hRetOrg electroporated at D42 with pCAG::GFP at 15V.**

**Supplementary Movie S3. D45 hRetOrg electroporated at D42 with pCAG::GFP at 40V.**

**Supplementary Movie S4. D45 hRetOrg electroporated at D42 with pCAG::GFP at 65V.**

**Supplementary Movie S5. Time-lapse of whole hRetOrg electroproated with pCAG::GFP**

**Supplementary Movie S6. Live tracking of mitochondria in D45 hRetOrg electroporated with pEF1α::mitoGFP**.

## Acknowledgements

We thank Connie Cepko for providing the pCAG::GFP and pUbC::mCherry constructs, and Kevin Eade and his team at the LMRI for their advice and training on organoid culture techniques. S.d.S. was supported by NEI R01EY033385, the ARVO Foundation for Eye Research (ARVO/Genentech AMD Research Fellowship Grant), and the Knights Templar Eye Foundation. We also acknowledge support from NIH CORE Grant P30 EY08098 and an unrestricted grant from Research to Prevent Blindness to the Department of Ophthalmology.

R.T.P. was supported by NIMH 5R01MH124695 and 1R21MH132015.

## Notes

### Competing Interest Statement

The authors have declared no competing interest.

